# Complex optogenetic spatial patterning with split recombinase

**DOI:** 10.1101/2024.11.07.622567

**Authors:** Ian S. Kinstlinger, Erin M. Neu, Shea K. Mowry, Cristina Tous, Darrell N. Kotton, Wilson W. Wong

**Affiliations:** Department of Biomedical Engineering and Biological Design Center, Boston University, Boston, MA 02215, USA; Center for Regenerative Medicine, Boston University and Boston Medical Center, Boston, MA 02118, USA; The Pulmonary Center and Department of Medicine, Boston University School of Medicine, Boston, MA 02118, USA

## Abstract

Light is a powerful and flexible input into engineered biological systems and is particularly well-suited for spatially controlling genetic circuits. While many light-responsive molecular effectors have been developed, there remains a gap in the feasibility of using them to spatially define cell fate. We addressed this problem by employing recombinase as a sensitive light-switchable circuit element which can permanently program cell fate in response to transient illumination. We show that by combining recombinase switches with hardware for precise spatial illumination, large scale heterogeneous populations of cells can be generated *in situ* with high resolution. We envision that this approach will enable new types of multicellular synthetic circuit engineering where the role of initial cell patterning can be directly studied with both high throughput and tight control.

## Introduction

Tissue morphogenesis during development arises from the exquisite coordination of cell proliferation, migration, and differentiation in response to cues from surrounding cells or from the environment. In particular, paracrine signaling through soluble morphogens is responsible for long-range cell-cell communication and represents a conserved mechanism for encoding and transmitting positional information.^1^

Across a variety of well-studied patterning systems, morphogen production is not homogenous within a tissue, but rather is concentrated in spatially defined signaling centers. For example, a signaling center in the posterior presomitic mesoderm serves as localized source of Wnt3a ligand, which forms a posterior-anterior gradient and drives somite formation.^2^ During lung development, branching morphogenesis is driven not by a continuum of Fgf10-secreting mesenchyme cells, but by localized centers of high expression.^3^ In these examples, morphogenesis is further mediated by subsequent soluble factors produced in response to the initial morphogen. Thus, the form and function of developing tissues may be encoded in the coupling between the spatial organization of the signaling center and the response of the downstream signaling network.

Synthetic development – the application of forward-engineering principles to the study of developmental biology – represents a promising avenue towards unraveling the molecular basis of morphogenesis.^4^ By abstracting naturally occurring signaling networks away from their native contexts, a more general and comprehensive understanding of their design principles can be gained.^5–8^ More recently, completely synthetic morphogen gradients have been demonstrated and used to pattern discrete domains of engineered cells^9^.

The repertoire of tools for synthetic development currently lacks a flexible and scalable strategy to spatially pattern cell fates. In the examples discussed above, only very rudimentary initial geometries of signaling centers (“sender cells”) and surrounding cells (“receiver cells”) were achieved, by seeding cells into a physical mold. Elsewhere in the synthetic biology literature, light-responsive proteins have been exploited to perturb developing systems with precise spatial control.^10–12^ While optical stimuli are evidently well-suited for spatial control, existing approaches are neither compatible with large-scale or high-throughput patterning, nor are they sufficiently modular to integrate alongside engineered genetic circuits to spatially define fully customized cell phenotypes.

Here, we employ light-inducible split recombinase as an effector to irreversibly transduce patterned optical inputs into engineered transcriptional outputs (i.e. cell fates). We describe an economical (∼$1200 USD) illumination system which can flexibly pattern large areas (∼100 cm^2^) with high resolution (∼250 μm) using a short exposure duration (≤5 min). We optimize and characterize the performance of this system and demonstrate high-fidelity patterning across entire multi-well plate areas using digital photomasks. The capacity to scalably and flexibly assign cell phenotype using patterned light enables a new generation of synthetic developmental biology where the spatial organization of cell populations becomes a tunable variable in the experimental design.

## Results

### Patterned stimulation of split recombinase

Our group previously established a library of high-performance inducible split recombinases, including split Cre and Flp fused to light inducible dimerization domains^13^. Because several of these split Cre and Flp proteins showed low basal activation and high dynamic range upon uniform exposure to blue light, we hypothesized that they would be able to reproduce a spatially heterogeneous optical pattern as a pattern of reporter expression in mammalian cells. We further envisioned that optogenetic patterning could be utilized to specify domains of morphogen sender and receiver cells (**Fig. 1a**), where the resulting cell phenotypes could be flexibly programmed through recombinase-actuated reporters (**Fig. 1b**).

**Fig. 1:**
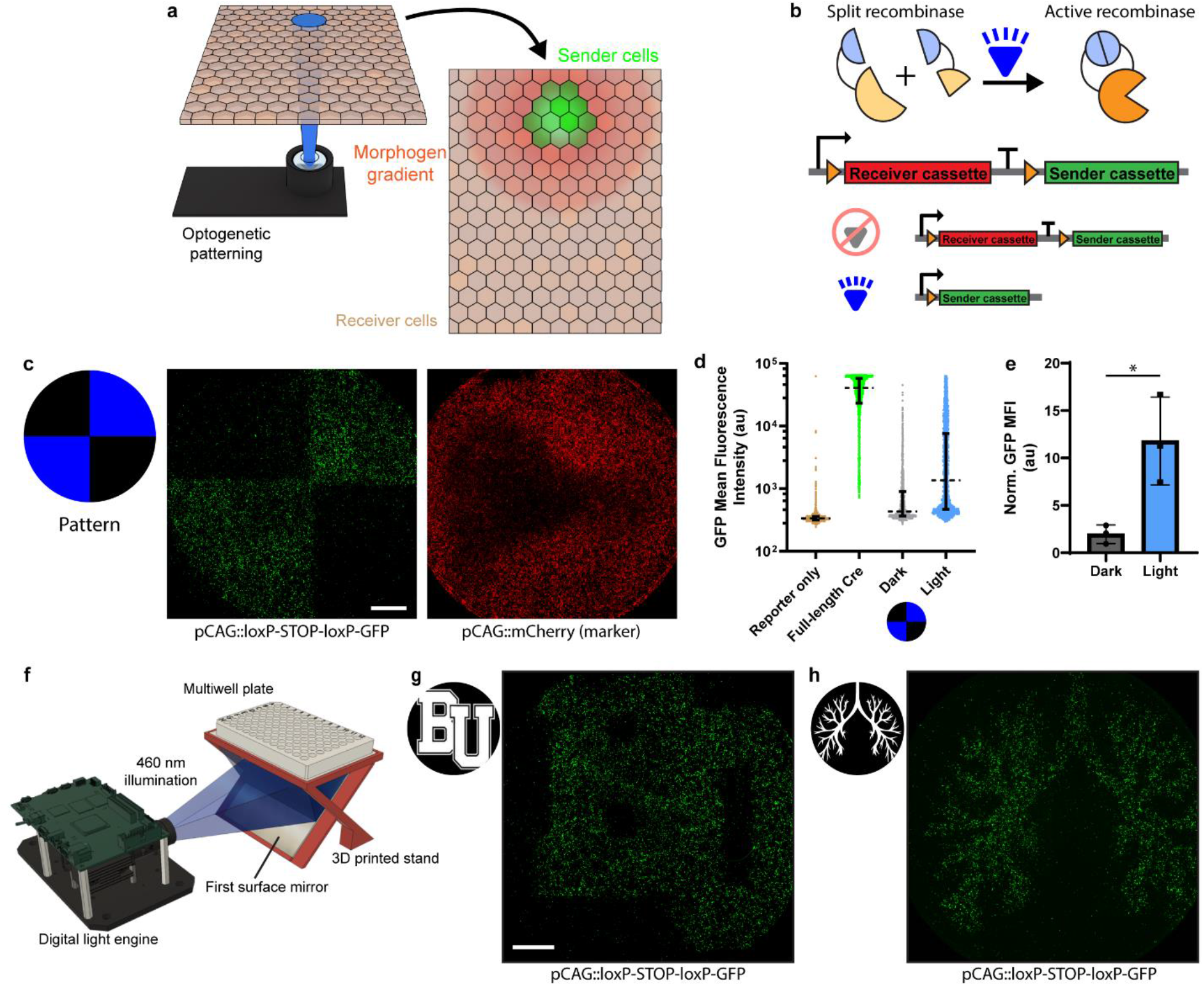
Split recombinase transduces heterogeneous illumination into cell fate patterns. **a)** Envisioned application of spatial optogenetic patterning to define populations of morphogen sender/receiver cells. **b)** Schematic of blue light-inducible split recombinase and reporter scheme for sender/receiver cell specification. **c)** A quadrant pattern of blue light was projected onto transiently transfected HEK293T cells. Epifluorescence microscopy shows faithful reproduction of the illumination pattern in the GFP Cre reporter channel, with transfected cells across all areas of the well. Scale bar = 2 mm. **d)** Distributions of GFP intensity within illuminated and dark pattern regions as well as controls. Each dot represents one transfected cell (as identified by binary image segmentation); dashed line shows median±i.q.r from n=1 representative biological replicate. **e)** Comparison of GFP signal between light and dark pattern regions; data are presented as mean±s.d. for N=3 biological replicates. * indicates p<0.05. **f)** Schematic of the well plate illumination apparatus used throughout the manuscript. **g,h)** Representative epifluorescence micrographs of patterns in transiently transfected HEK293T cells; pattern designs shown at left Scale bar = 2mm. The design in **(h)** is derived from a stylized illustration of the lung by Alexander Mikhailov, used under standard license from iStock.

We selected the split recombinase which showed the highest fold change upon blue light induction (iCre_N-229_-pMag + nMag-iCre_230-C_) and tested its performance using transient transfection in HEK293T cells. Throughout this manuscript, we employed a Cre reporter which consists of GFP downstream of a loxP-flanked transcriptional terminator, and we co-transfected a constitutive transfection marker. We observed that patterned illumination with 460 nm light yielded well-defined domains of cells with and without reporter activation, separated by a sharp boundary (**Fig. 1c,d**). Further image analysis showed that approximately 40% of cells within the illuminated part of the pattern expressed reporter, compared with approximately 10% within the dark region (**Fig. 1e**). Because <1% of cells in reporter-only controls were GFP^+^, we concluded that split Cre recombination in the dark is the major source of noise in the system, as opposed to reporter read-through. Even in GFP^+^ cells, the intensity of GFP expression driven by split Cre was substantially lower than observed with full-length Cre, in agreement with our previous findings^13^.

In order to establish a workflow to reproducibly stimulate cells with light, we developed an illumination apparatus to accommodate standard multiwell plates (**Fig. 1f**). The apparatus consists of a high-performance digital light engine, an optical-grade first surface mirror, and a custom 3D printed plate and mirror stand. These components are mounted on T-slot aluminum extrusion rails so that the working distance from projector to sample can be continuously adjusted. The light engine projects standard image files as digital photomasks and has an adjustable LED current value which controls the optical power. With this hardware, we evaluated our capacity to pattern increasingly complex designs and concluded that this combination of optics and split recombinase was amenable to rapid (5 min) induction of high-fidelity patterns in diverse shapes (**Fig. 1g,h**). However, we also noted that cells within the pattern were fairly sparse and often unevenly seeded, and that undesired reporter expression in the dark regions of the patterns detracted from the overall contrast.

### Optimization of cell seeding and illumination parameters

Achieving high-fidelity optogenetic patterns is dependent on both efficient cell transfection and high cell density. However, because standard transfection protocols perform optimally at intermediate cell density, these goals appeared mutually exclusive. For example, our early experiments employed optimized transfection conditions and yielded relatively sparse cell patterns (**Fig. 1g,h**). We overcame this hurdle by decoupling transfection from cell patterning (**Fig. 2a**). We found that by transfecting cells at intermediate density (80×10^3^ cell cm^-2^), then subsequently re-plating them at high-density (325×10^3^ cell cm^-2^), we could generate dense monolayers of efficiently transfected cells. At this stage, we also switched from a completely transient transfection protocol to a stably integrated line expressing the split Cre, with only a transient reporter (see Methods). Patterned illumination of these monolayers yielded on-regions with higher GFP^+^ cell density and crisper boundaries between on- and off-regions (**Fig. 2b,c**).

**Fig. 2:**
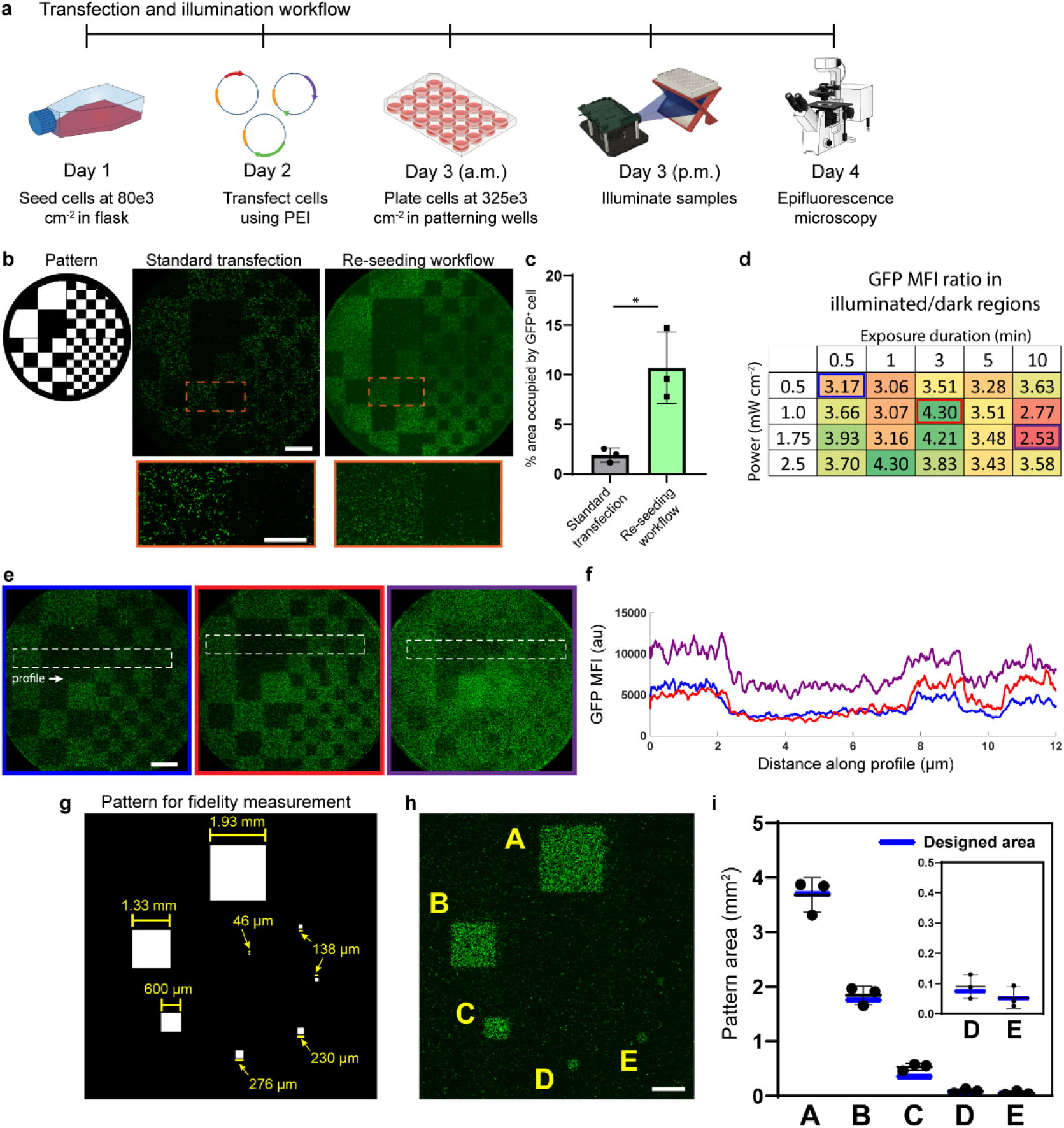
Optimized seeding and illumination workflows yield high-contrast, high-resolution patterns. **a)** Timeline depicting the experimental workflow for cell seeding, transfection, and illumination. Well plate and microscope icons are by DBCLS, used under CC-BY. **b)** Side-by-side comparison of the same design patterned with a standard transfection protocol versus the workflow schematized in **(a)**. Note that the re-seeding workflow includes also the switch from transient transfection to an integrated split Cre-expressing line (see Methods). Scale bar = 2 mm; 1 mm in zoom-in region **c)** Area coverage was computed from binarizing the GFP channel in on-regions of the quadrant pattern shown in **Fig. 1c** (standard transfection) and the largest square of the pattern shown in **(g)** (re-seeding workflow). N=3 biological replicates; data are plotted as mean±s.d.; * indicates p<0.05. **d)** On- and off-region reporter intensity was quantified and ratio computed from epifluorescence images of wells patterned with various power and duration settings. N = 1 biological replicate. **e)** Epifluorescence micrographs corresponding to the boxed settings in **(d)**. Scale bar = 2mm. **f)** Intensity profile along the indicated region in **(e)** representing the mean value at each horizontal position. Profiles were smoothed with a moving-average function. **g)** Pattern design for quantifying dimensional accuracy with nominal feature dimensions. **h)** Epifluorescence micrograph of the dimensional accuracy pattern, scale bar = 2 mm. **i)** Actual feature areas were computationally extracted (see Methods) and compared to the nominal dimensions depicted in **(g)** (blue lines). N = 3 biological replicates; data are plotted as mean±s.d.

In order to optimize the contrast between on- and off-regions, we varied the optical power and exposure duration during blue light illumination. Crucially, we observed that while the off regions of our digital photomasks have a pixel intensity value of zero, the LED array in the light engine generates roughly 0.04 mW cm^-2^ of optical power (roughly one order of magnitude below typical patterning conditions for on-regions). We therefore expected that patterning quality would decline with prolonged exposure due to cumulative basal photon flux through the off-pixels. Indeed, we found that exposure times exceeding 3 minutes, and especially exceeding 5 minutes, led to reduced pattern contrast due to higher intensity reporter expression in the off-regions (**Fig. 2d**). Moreover, we found that patterning times as short as 1-3 min were sufficient to activate reporter in the on-regions with minimal expression in the off-regions. Side-by-side comparison of the patterns produced by the optimal illumination condition (**Fig. 2e**, middle) and two sub-optimal conditions highlights losses in pattern quality from incomplete activation of the on regions (**Fig. 2e**, left) and excessive noise caused by overexposure of the off regions (**Fig. 2e**, right). Plotting the profile of GFP intensity across on- and off-region boundaries indicates that the boundary sharpness is not dramatically affected by the choice of illumination intensity, but again highlights the substantial increase in off-region basal reporter level (**Fig. 2f**).

### Patterning resolution and dimensional accuracy

Before attempting to determine the resolution of this system, we maximized resolution at the hardware level by minimizing the working distance between the light engine and sample and found that each pixel of a 1920×1080 px image corresponded to 46 μm. We characterized the resolution in this configuration by projecting a pattern of square regions of decreasing size, down to exactly 1 pixel (**Fig. 2g**). Qualitative inspection of the images suggested that the largest five squares (1.93 mm side length down to 230 μm) could be patterned and retain their designed shape and size (**Fig. 2h**). Below this size, the signal in the expected position of the squares became difficult to distinguish from the speckled background signal typical of these patterning experiments.

In order to make an unbiased evaluation of the resolution limit and quantify the areas of the patterned features, we developed an image processing algorithm to convert the raw epifluorescence images into binary feature maps. We found that conventional thresholding was poorly ineffective for segmenting these multicellular features due to the lack of continuity between cells in the on-regions. Instead, we turned to a 2D convolution algorithm which assigns each pixel in the image a value based on the intensity of pixels in a surrounding neighborhood, specified by a user-defined kernel. Convolution was successful at segmenting the patterned features because it returns large values for pixels in on-regions (i.e. with many high-MFI neighbors) and small values for pixels in off-regions. We post-processed the binary feature maps generated via 2D convolution to measure the area of each feature.

We found that squares down to side length 230 μm (5 px) could be reproducibly patterned with high dimensional accuracy (**Fig. 2h,i**). The next smallest feature (138 μm, 3 px) was detected in one biological replicate, but could not be consistently identified and was thus considered below the resolution limit. The measured area of almost all features fell within ±10% of the designed area. Notably, the measured area of the features typically slightly exceeded the designed area.

### Large-scale pattern design and realization

Two key design goals for this study were to enable flexible and on-demand patterning of different designs within each well of a multiwell plate, and to push the achievable limit of length scale for high-resolution patterning. Towards the first goal, we developed an image processing chain in MATLAB which takes in individual image files or sets of images and outputs a single 1920×1080 px grayscale image such that the projected patterns align with the wells of a multiwell plate (**Fig. 3a**). Simultaneous patterning across an entire well plate requires a longer working distance from the light engine to the sample, such that each pixel covers 83×83 μm. We used this workflow to pattern a set of related but distinct patterns across a 24-well plate using a single 5 min exposure (**Fig. 3b**).

**Fig. 3:**
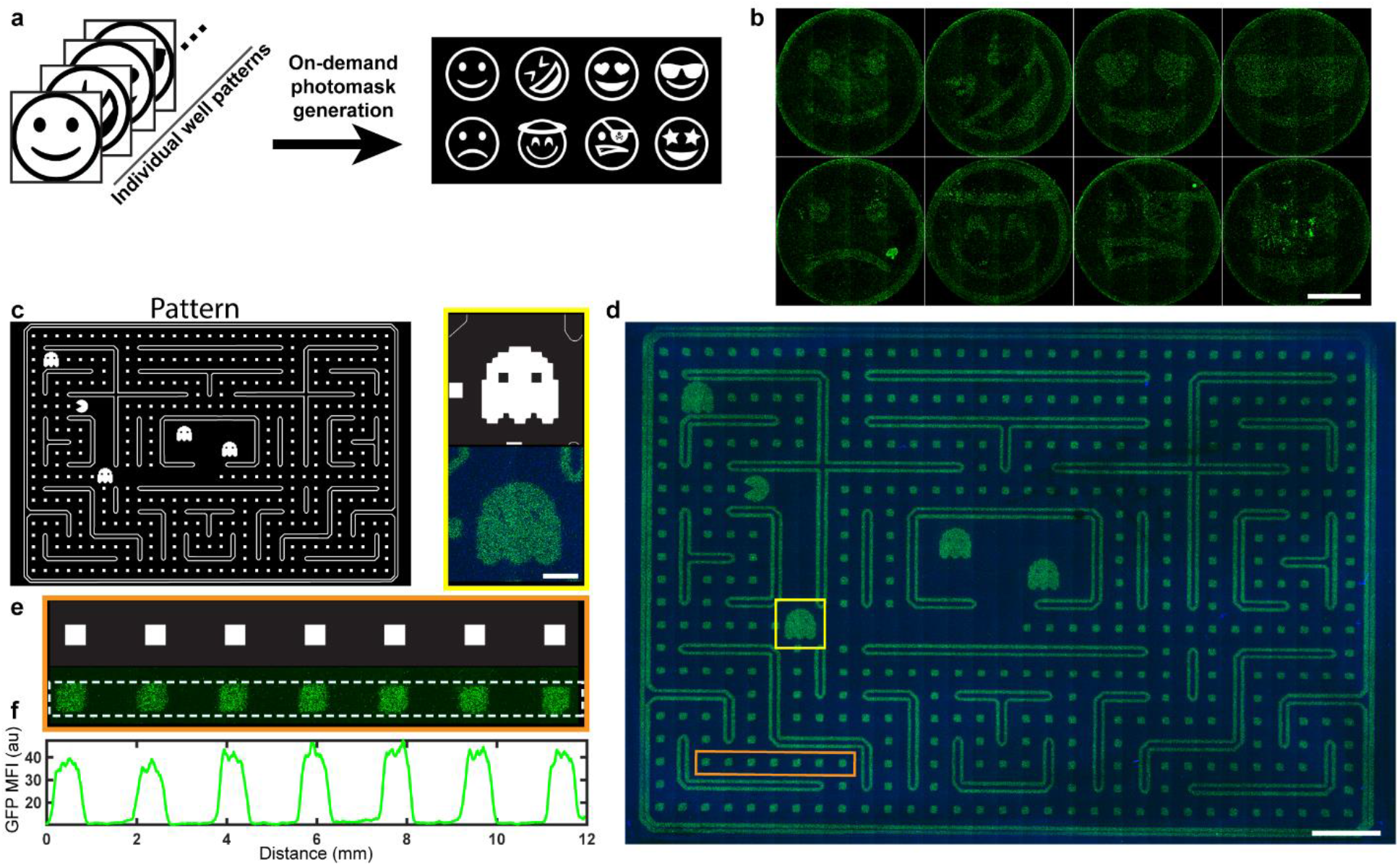
Split recombinase optogenetics enables programming of large-scale spatial patterns. **a)** Workflow schematic for the production of digital photomasks for arbitrary image inputs, demonstrated here with a set of emojis (input images are Adobe stock images, used under a standard license). **b)** Montage of patterns from the design depicted in **(a)**. Images were processed for ease of visualization with Gamma correction (factor = 1.5) and 2×2 binning; scale bar = 5 mm. **c)** A pattern reminiscent of the Pac-Man arcade game was designed in Adobe Illustrator to demonstrate large area optogenetic patterning. **d)** Stitched epifluorescence micrographs of cells illuminated with the pattern in **(c)**. The blue channel is pCAG::BFP which was transfected as a marker. Scale bar = 1 cm. Yellow inset scale bar = 2 mm. **e)** Epifluorescence micrographs of the boxed orange region of **(d). f)** Intensity profile along the indicated region in **(e)** representing the mean value at each horizontal position. Profile was smoothed with a moving-average function.

Finally, we sought to produce a single coherent pattern across the footprint of an entire multiwell plate as a proof point for spatial optogenetics at the length scale of multiple centimeters. We adapted our MATLAB tool to align a single pattern across an entire 1-well plate and performed a single 5 min exposure (**Fig. 3c**). We found that cells were dense and homogeneous across this 10×7 cm pattern and that the entire design was faithfully transferred to the cells without significant aberrations or artifacts (**Fig. 3d**). Features as small as 2-4 pixels are visible in the final pattern, and features on the order of 1 mm were dimensionally accurate and marked by sharp boundaries (**Fig. 3e**).

## Discussion

In this paper, we demonstrated the use of light inducible split-recombinase as an effector for large-scale spatial patterning of cell phenotype. We demonstrated that programming of cell fate through a digitally designed pattern can be performed at relatively high resolution (<250 μm features) and across expansive patterning areas. We identified trends in the effects of patterning parameters on pattern contrast and fidelity towards the realization of dense patterns with low background noise. Our data also highlight that spatial optogenetic patterning can be a high-throughout proposition, with optimal patterning times under 5 min and the ability to pattern distinct patterns across different wells simultaneously. Moreover, recombinase is a highly modular effector and can, in principle, be used to drive a permanent phenotype switch for a wide range of potential outputs. Logic gates and biocomputation architecture have already been developed for recombinases,^14^ suggesting that it could be possible to use optogenetic patterns as inputs into logic computation circuits.

While optogenetic patterning^15^ and coupling of those patterns to downstream biocomputing circuits^16^ were achieved in engineered bacteria well over a decade ago, it has been more challenging to realize equivalent patterns in mammalian cells. Undoubtedly, optogenetic approaches have had tremendous influence as tools for biological discovery and engineering; however, many such studies either do not deliver light in a spatially-defined manner, or perform spatial patterning at length scales on the order of tens of microns, typically through the optical train of a microscope.^12,17^ The small number of published studies that investigate mammalian optogenetic patterning across larger areas have generally relied on physical photomasks,^18,19^ which may be time- and/or labor-intensive to produce and are not ideally suited for studies where the experimenter wishes to flexibly iterate through many pattern geometries. We therefore believe that our digital photomask-based approach may represent a valuable tool for the community and empower studies examining the role of initial patterns of gene expression in the development of multicellular morphologies.

The patterns presented in this work represent, to our knowledge, the largest areas optogenetically patterned at sub-millimeter resolution. It is reasonable to ask whether there is a clear benefit to patterning in this size regime, especially when so many developmental biology models are orders of magnitude smaller. While the field is in its infancy, several published studies of mammalian cell patterning through synthetic paracrine interactions do produce patterns on the order of several millimeters^8,9,20^ Controlling the initial placement of sender and receiver cells in such systems *in situ* with high throughout could be a good use case for this form of patterning. Separately, we note that in the field of biomaterials, light has been harnessed for spatially controlling polymer chemistry at length scales on the order of tens of microns^21^ all the way up to tens of millimeters^22^. The former are useful for exquisitely manipulating the mechanical microenvironment around individual cells, whereas the latter represent a promising strategy towards engineering functional tissues. Perhaps, as mammalian synthetic biology develops and matures, we will find that optogenetic patterning tools spanning the same range of length scales can likewise be harnessed for both fundamental biological investigation and for fabricating functional and programmable multicellular assemblies.

## Methods

### Plasmid design and assembly

Constructs were designed through standard molecular biology protocols and Gibson homology-based assembly cloning. Initial transient expression plasmids for the split blue-light inducible Cre were reported in Ref ^13^. All sequences were verified by sequencing prior to use in mammalian expression. The full plasmid maps of all constructs will be made available at the time of publication.

### Mammalian cell culture and transient transfection

HEK293T cells were maintained at 37 °C, 5% CO_2_ in DMEM (Corning) supplemented with 5% Fetal Bovine Serum (FBS; Gibco) and 1% penicillin-streptomycin (Thermo Fisher). For standard transient transfection experiments, cells were resuspended to a density of 250×10^3^ cells ml^-1^ and seeded at a volume of 100μl in 96-well plates or 500μl in 24-well plates. For illumination experiments, black-walled 96-well (Nunc) and 24-well (Ibidi) plates were used to reduce light spillover between wells. We also supplemented the cell culture media with 2mM tartrazine (Chem Impex), a biocompatible dye which we hypothesized could attenuate background illumination based on its previous use as a photoabsorber.^22^ While we ultimately did not conclude that the addition of the dye had a statistically significant effect on pattern quality, we included it in replicate experiments for consistency.

Transient transfection was performed using polyethylenimine (PEI) (Polysciences, 23966, prepared as a .323 g l^-1^ stock). For each transfected well of a 24-well plate, 500ng DNA was diluted in 25 μl 0.15M NaCl and vortexed prior to the addition of 25 μL PEI, diluted to 16% v/v in 0.15M NaCl. DNA/PEI mixes were vortexed again and incubated for 10-15 min at room temperature prior to addition to cells. In some experiments, cells were transfected the day after seeding while in others, DNA was added at the same time the cells were seeded (reverse transfection). In transient patterning experiments, the ratio of DNA components was 1 part N-terminal Cre split:1 part C-terminal Cre split:1 part transfection marker: 13 parts Cre reporter. For reporter only negative controls, the Cre splits were replaced by a dummy plasmid (CAG promoter with no coding sequence) and for wild-type Cre positive controls, the Cre splits were replaced by the full-length Cre sequence driven by a CAG promoter. Transfection marker was either mCherry or BFP (for microscopy experiments) or iRFP-720 (for flow cytometry), driven by a CAG promoter. Plates were wrapped in foil to prevent undesired activation by ambient light.

### Stably integrated drug-inducible split recombinase line

While our initial experiments to validate optogenetic recombinase patterning were performed with transiently transfected components, we sought to develop a stable line of HEK293T cells with which to further optimize and evaluate this system. We reasoned that a stable line could provide greater reproducibility across experiments and serve as an “off-the-shelf” cell line for building more complex circuits downstream of the recombinase. Because our initial experiments showed moderate levels of dark activation, we chose to drive split Cre expression with an inducible tet promoter, such that the system would only be “unlocked” for patterning by the addition of doxycycline.

Non-targeted integration was performed in HEK293T cells using hyperactive piggyBac transposase^23^ and PEI transfection as described above. We simultaneously integrated three plasmids, each constructed in a piggyBac backbone: pTre::iCre_N-229_-pMag, pTre::nMag-iCre_230-C_, and pEF1α-rtTA. DNA was mixed in a ratio of 1 part piggyBac transposase:1.67 parts each target plasmid. Following integration, cells were expanded and selected with puromycin (2 μg/mL; Invivogen) for 10 days. Blue light-responsive cells were purified by Fluorescence-Activated Cell Sorting (FACS) and cultured in complete medium supplemented with 1 ng μl^-1^ doxycycline (dox) as a measure to prevent gene silencing.^24^

### Illumination hardware

Blue light illumination was performed using a DLP LightCrafter 4500 digital light engine with independently addressable red, green, and blue LED arrays (EKB Technologies, E4500MKIIRGB). The display image was controlled via HDMI connection to a PC and the LED current was controlled over USB through the LightCrafter control software. To illuminate the bottom surface of a multiwell plate for adherent cell patterning, a mirror (First Surface Mirror, 1/8” thick) was positioned at a 45° angle to redirect the light upwards the plate. We designed and 3D printed a custom stand to hold the mirror and plate, and the light engine as well as the stand were fastened to a frame made from aluminum T-slot extrusion. For single-well patterning experiments, we lasercut a 14 mm hole in a sheet of ¼” black acrylic, which mounts atop the 3D printed stand. The black plastic blocks illumination of all wells except the one positioned above the hole. Optical power output from the light engine was measured using a USB power meter (Thorlabs, PM16-120).

### Digital photomask generation

Photomasks were generated with a custom MATLAB script. The size and spacing of individual wells in a 24-well plate was used to define the pixel coordinates that would project onto each well, with empirical adjustments made to achieve optical alignment; a similar process was repeated for the single-well patterning configuration. To generate a photomask from an arbitrary image, the image is read into MATLAB as an array, converted to grayscale and double format, and resized to match the pixel dimensions of the patterning area. In minor variations of the script, a loop can be used to tessellate a single image across multiple well positions in a plate, or to assign each layer of an image stack to a different well position. While the image arrays are rectangular, the physical wells are circular, so a masking operation is used to produce a circular pattern. The final photomask is exported as an image file and displayed on the digital light engine.

### Optimized transfection and patterning

For patterning experiments with the exception of Fig. 1, we employed the following workflow, as schematized in Fig. 2a. 6×10^6^ integrated and sorted HEK cells with dox-inducible expression of blue-light inducible Cre were seeded in a T75 flask in 15 ml culture medium. The following day, the PEI transfection procedure described above was scaled up for the T75 flask (19.5 μg DNA in 975 μl 0.15M NaCl, mixed with 975 μl of 16% v/v PEI). The DNA was split into 1 part transfection marker + 7 parts Cre reporter. One day later, the transfected cells were collected with trypsin and re-seeded in 24-well black-walled plates at a density of 1×10^6^ cells per well, and dox was added to the media (2 ng/μl). On the same day (at least 5 hours after seeding), the plate was illuminated using optimized settings for the appropriate optical configuration. In single-well patterning mode, each pattern was exposed for 3 min at a power density of 1.0 mW cm^-2^ and in whole-plate patterning mode, each pattern was exposed for 5 min at 1.0 mW cm^-2^. The day after illumination, patterning was evaluated using epifluorescence microscopy.

### Epifluorescence microscopy

Samples were imaged on a Cytation 5 cell imager (BioTek) equipped with LED illumination sources and filter cubes for detection of GFP, mCherry, and BFP. GFP used a 456 nm LED paired with a 469/525 nm (excitation/emission) filter set; mCherry used a 554 nm LED paired with a 556/600 nm filter set; BFP used a 365 nm LED paired with a 377/447 nm filter set. For each experiment, exposure time was set for the transfection marker and Cre reporter channels using the wild-type Cre positive control condition. To image 24-well plate samples, a 4×4 montage was collected using the 4x objective and to image an entire multiwell plate footprint, a 31×21 montage was collected. Tiles were stitched to produce the final image using a custom Python script.

### Quantitative image analysis

GFP mean fluorescence intensity (MFI) was quantified in on- and off-regions of patterns by drawing a rectangular ROI in Fiji and measuring the mean pixel intensity. GFP intensity of individual cells within a pattern (Fig. 1d) was determined by using the Analyze Particles function in Fiji to segment cells in the mCherry (transfection marker) channel. Each segmented cell was then treated as an ROI for measurement of mean pixel intensity in the GFP channel. Intensity profiles within images were extracted using the Plot Profile function in Fiji and were smoothed with a moving average function in MATLAB before plotting.

The area of patterned features was quantified in MATLAB by performing 2D convolution of the stitched GFP channel image (1992×1992 px) with a uniformly-valued kernel of dimension 149×149 px. The result of the convolution was morphologically dilated, then binarized and cleaned by automated removal of pixel regions smaller than a defined threshold. The position and area of the segmented features were extracted using the regionprops function in MATLAB.

## Acknowledgements

We thank all members of the Wong Lab for helpful discussions. I.S.K. acknowledges funding from a postdoctoral NRSA award from the NIH/NHLBI (F32HL160194), the TL1 Postdoctoral Fellowship in Regenerative Medicine (TL1TR1410), and a Kilachand postdoctoral fellowship from the Boston University Biological Design Center. D.N.K. and W.W.W. acknowledge funding from an Allen Distinguished Investigators award from the Allen Institute (12963).

## Notes

### Competing Interest Statement

The authors have declared no competing interest.

## References

1. Kicheva, A. & Briscoe, J. Control of Tissue Development by Morphogens. Annu. Rev. Cell Dev. Biol. 39, 91–121 (2023).

2. Aulehla, A. & Pourquie, O. Signaling Gradients during Paraxial Mesoderm Development. Cold Spring Harb. Perspect. Biol. 2, a000869–a000869 (2010).

3. Bellusci, S., Grindley, J., Emoto, H., Itoh, N. & Hogan, B. L. M. Fibroblast Growth Factor 10 (FGF10) and branching morphogenesis in the embryonic mouse lung. Development 124, 4867–4878 (1997).

4. Ebrahimkhani, M. R. & Ebisuya, M. Synthetic developmental biology: build and control multicellular systems. Curr. Opin. Chem. Biol. 52, 9–15 (2019).

5. Li, P. et al. Morphogen gradient reconstitution reveals Hedgehog pathway design principles. Science 360, 543–548 (2018).

6. Toda, S., Blauch, L. R., Tang, S. K. Y., Morsut, L. & Lim, W. A. Programming self-organizing multicellular structures with synthetic cell-cell signaling. Science eaat0271 (018) doi:10.1126/science.aat0271.

7. Stapornwongkul, K. S., de Gennes, M., Cocconi, L., Salbreux, G. & Vincent, J.-P. Patterning and growth control in vivo by an engineered GFP gradient. Science 370, 321–327 (2020).

8. Zhu, R. et al. Reconstitution of morphogen shuttling circuits. Sci. Adv. 9, eadf9336 (2023).

9. Toda, S. et al. Engineering synthetic morphogen systems that can program multicellular patterning. Science 370, 327–331 (2020).

10. Johnson, H. E., Djabrayan, N. J. V., Shvartsman, S. Y. & Toettcher, J. E. Optogenetic Rescue of a Patterning Mutant. Curr. Biol. 30, 3414-3424.e3 (2020).

11. Legnini, I. et al. Spatiotemporal, optogenetic control of gene expression in organoids. Nat. Methods 20, 1544–1552 (2023).

12. McNamara, H. M. et al. Optogenetic control of Nodal signaling patterns. Preprint at 10.1101/2024.04.11.588875 (2024).

13. Weinberg, B. H. et al. High-performance chemical- and light-inducible recombinases in mammalian cells and mice. Nat. Commun. 10, 4845 (2019).

14. Weinberg, B. H. et al. Large-scale design of robust genetic circuits with multiple inputs and outputs for mammalian cells. Nat. Biotechnol. 35, 453–462 (2017).

15. Levskaya, A. et al. Engineering Escherichia coli to see light. Nature 438, 441–442 (2005).

16. Tabor, J. J. et al. A Synthetic Genetic Edge Detection Program. Cell 137, 1272–1281 (2009).

17. Martínez-Ara, G. et al. Optogenetic Control of Apical Constriction Induces Synthetic Morphogenesis in Mammalian Tissues. http://biorxiv.org/lookup/doi/10.1101/2021.04.20.440475 (2021) xdoi:10.1101/2021.04.20.440475.

18. Repina, N. A. et al. Engineered Illumination Devices for Optogenetic Control of Cellular Signaling Dynamics. Cell Rep. 31, 107737 (2020).

19. Polstein, L. R., Juhas, M., Hanna, G., Bursac, N. & Gersbach, C. A. An Engineered Optogenetic Switch for Spatiotemporal Control of Gene Expression, Cell Differentiation, and Tissue Morphogenesis. ACS Synth. Biol. 6, 2003–2013 (2017).

20. Zhang, X. et al. Post-Transcriptional Modular Synthetic Receptors. Preprint at 10.1101/2024.05.03.592453 (2024).

21. Shadish, J. A., Benuska, G. M. & DeForest, C. A. Bioactive site-specifically modified proteins for 4D patterning of gel biomaterials. Nat. Mater. 18, 1005–1014 (2019).

22. Grigoryan, B. et al. Multivascular networks and functional intravascular topologies within biocompatible hydrogels. Science 364, 458–464 (2019).

23. Yusa, K., Zhou, L., Li, M. A., Bradley, A. & Craig, N. L. A hyperactive piggyBac transposase for mammalian applications. Proc. Natl. Acad. Sci. U. S. A. 108, 1531–1536 (2011).

24. DiAndreth, B., Wauford, N., Hu, E., Palacios, S. & Weiss, R. PERSIST platform provides programmable RNA regulation using CRISPR endoRNases. Nat. Commun. 13, 2582 (2022).

